# The three-dimensionally articulated oral apparatus of a Devonian heterostracan sheds light on feeding in Palaeozoic jawless fishes

**DOI:** 10.1101/2023.08.22.554283

**Authors:** Richard Dearden, Andy Jones, Sam Giles, Agnese Lanzetti, Madleen Grohganz, Zerina Johanson, Stephan Lautenschlager, Emma Randle, Philip C. J. Donoghue, Ivan J. Sansom

## Abstract

Attempts to explain the origin and diversification of vertebrates have commonly invoked the evolution of feeding ecology, contrasting the passive suspension feeding of invertebrate chordates and larval lampreys with active predation in living jawed vertebrates. Of the extinct jawless vertebrates that phylogenetically intercalate these living groups, the feeding apparatus is preserved only in the early diverging stem-gnathostome heterostracans and its anatomy remains poorly understood. Here we use X-ray microtomography to characterise the feeding apparatus of the pteraspid heterostracan *Rhinopteraspis dunensis* (Roemer, 1855). We show that the apparatus is composed of thirteen plates arranged approximately bilaterally, the majority of which articulate from the postoral plate. Our reconstruction of the apparatus shows that the oral plates would have been capable of movement within the dorso-ventral plane, but their degree of movement was limited. The functional morphology of the apparatus in *Rhinopteraspis* precludes all proposed interpretations of feeding except for suspension/deposit feeding and we interpret the apparatus as having served primarily to moderate the oral gape. This is consistent with evidence that at least some early jawless gnathostomes were suspension feeders and runs contrary to macroecological scenarios that envisage early vertebrate evolution as characterised by a directional trend towards increasingly active food acquisition.

## 1. Introduction

Feeding figures prominently in attempts to understand the evolutionary origins of vertebrates (Janvier 1996, Anderson *et al*. 2011). In contrast to invertebrate chordates, which exclusively suspension feed with either a ciliated pharynx or a mucus net (1), the dorso-ventrally closing jaws of living jawed vertebrates (crown-gnathostomes plus ‘placoderms’) (2–4) or antero-posteriorly moving system of cartilages (in cyclostomes, hagfishes and lampreys) (5–8) allow for a far broader range of feeding strategies. The evolution of these unique vertebrate feeding modes plays a major role in attempts to explain the evolution of vertebrate anatomy and the origins of its modern diversity (9,10). Prominently, the New Head Hypothesis (11–15), argues that the shift from suspension feeding to predation accompanied the emergence of neural crest, neurogenic placodes, and the accompanying evolution of a prechordal head. This hypothesis predicts the earliest vertebrates with a prechordal head, i.e. parts formed from trabecular elements of the neurocranium anterior to the notochord(16), had a predatory feeding mode.

The fossil record of Palaeozoic vertebrates provides a means of testing such hypotheses, but current interpretations of that record are far from decisive. In particular, heterostracans, an extinct group of jawless stem-gnathostomes, have been the focus of much of the debate over feeding in early vertebrates. This is because their oral region is more commonly and completely preserved than in any other such group, and they are often interpreted as one of the earliest diverging lineages of stem-gnathostomes. As such, heterostracans have the potential to inform on the feeding ecology of the earliest members of the gnathostome lineage (17). The heterostracan feeding apparatus is best known in pteraspids, where the oral region is characterised by distinctive macromeric dermal plates (18–22). The function of these plates has been much debated, variously interpreted as biting (19) or slicing (23) ‘jaws’, a cyclostome-like feeding apparatus (24–27), a sediment scoop (22), or a filtering structure (28–32). Equally varied are the inferred ecologies, with heterostracans interpreted as active predators (19), macrophagous selective predators (12,33), hagfish-like scavengers (24–27), herbivores (23), detritivores (22,34), or suspension feeders (including filter feeding) (10,20,35– 38). The most recent investigation suggests that heterostracans were suspension feeders because the analysed oral plates exhibited no evidence of the wear anticipated of a ‘tooth-like’ function (20).

The difficulty in studying the articulated heterostracan oral apparatus *in situ* contributes to this lack of consensus. In the rare cases where they are preserved, articulated heterostracan oral apparatuses consist of small plates suspended in encasing sediment. As a result, previous reconstructions of the oral apparatus have focused either on the gross arrangement of the apparatus, in which the morphology and detailed arrangement of the individual plates are not characterised (22,24,26,27), or describe isolated elements with little or no reference to articulated apparatuses (22,39,40). Evidently, the feeding ecology of heterostracans remains in its infancy and so we sought to advance understanding through a detailed characterization and reconstruction of the heterostracan oral apparatus. We used X-Ray microtomography to characterise the three-dimensionally (3D) articulated oral apparatus of an exceptionally well-preserved specimen of the pteraspid heterostracan *Rhinopteraspis dunensis* (Roemer, 1855). Using computed tomography, we generated volumetric models of the components of the oral apparatus and used these models to reconstruct their three dimensional arrangement in vivo. We use this reconstruction to assess competing hypotheses of heterostracan feeding.

## 2. Materials & Methods

### (a) Specimens and locality

*Rhinopteraspis dunensis* (Roemer, 1855) NHMUK PV P 73217 is housed in the collections of the Natural History Museum, London (NHMUK). In the museum catalogue the specimen is listed as being collected from “Odenspiel Quarry”, likely corresponding to Jaeger Steinbruch, a quarry near the village of Odenspiel, Reichshof, North-Rhine Westphalia, Germany or, possibly, outcrops in the local area (41,42). The Jaeger Quarry and surrounding outcrops expose sandstones and mudstones belonging to the Siegenian (?upper Pragian or lower Emsian, Lower Devonian) Odenspiel Formation (43,44), deposited on the northern margin of the Rhenohercynian Basin, which was a marginal transgressive and regressive delta-dominated setting (45). The Odenspiel Formation falls within the ‘Pararhenotypics subfacies’ of Jansen (46), representing a marginal marine, intertidal lagoonal setting preserving a restricted fauna of fish, bivalve molluscs, lingulid and terebratulid brachiopods, and eurypterids (41,42,47–49).

### (b) Terminology

Various terminologies have been applied to the pteraspid oral region. Here we follow the terminology of Randle and Sansom (50) and, where that is not possible, that of Blieck (51). We extend existing terminology to describe the nature and arrangement of the oral plates (Fig. 1G). For the anatomical axes of the animal as a whole we use dorsal/ventral in the dorsoventral axis, rostral/caudal for the sagittal axis, and dextral/sinistral for the transverse axis. When describing the oral apparatus itself we also use lateral/medial to describe lateral positions relative to the sagittal axis, ad/aboral to describe the surfaces of plates relative to the mouth (i.e. adoral is the surface of a plate facing the mouth, aboral the surface facing away), and proximal/distal to refer to positions on the oral plates relative to their articulation with the postoral plate with oral plate tips being distal.

**Figure 1.**
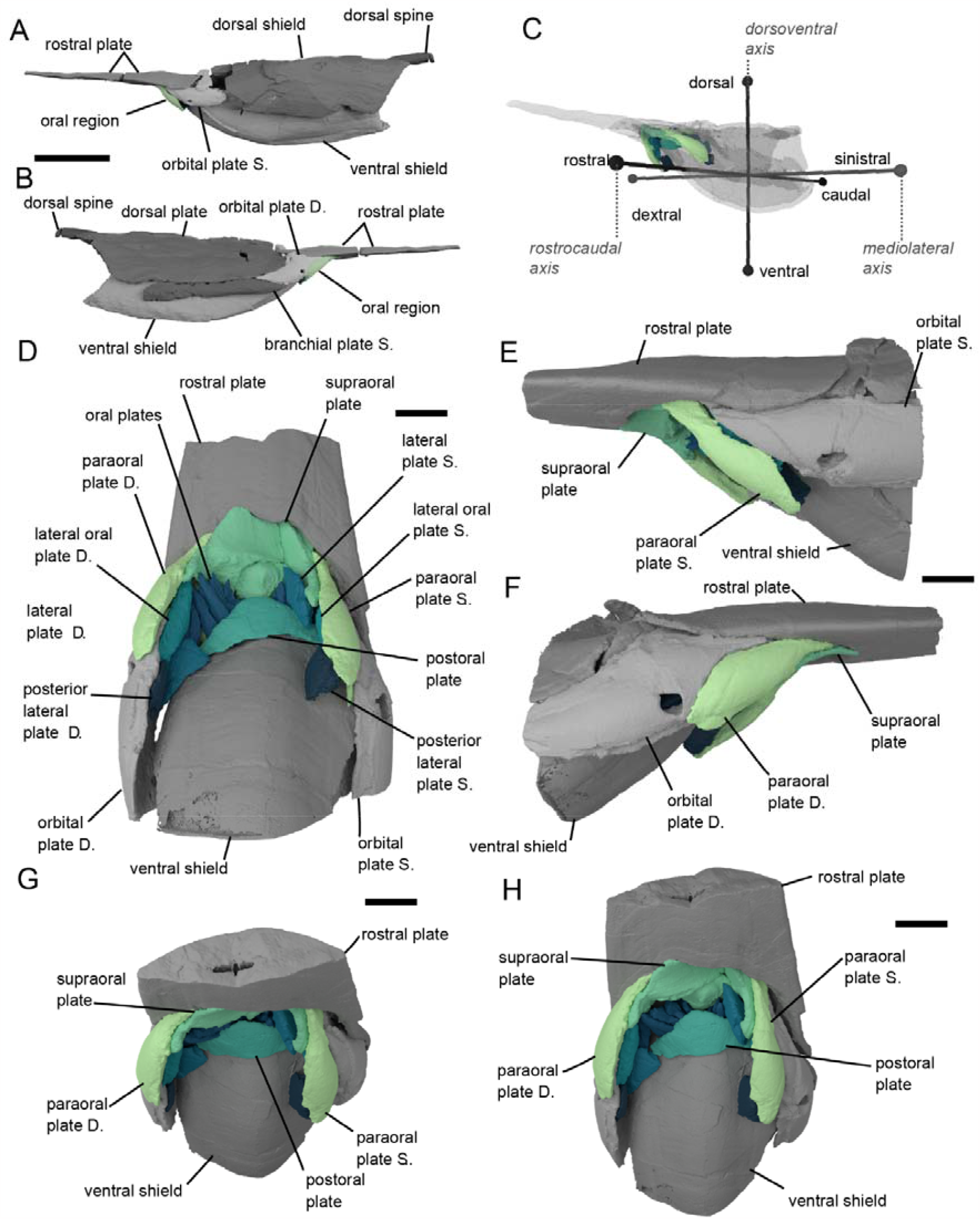
*Rhinopteraspis dunensis* NHMUK PV P 73217. A-C, Rendering of head shield based on computed tomographic data in (A) sinistral, (B) dextral view, and (C) transparent with scheme of anatomical axes. D-H, renders based on higher resolution data showing the oral apparatus in more detail in D, ventral view; E, sinistral view; F, dextral view; G rostral view; H, Rostro-ventral view. Green and blue parts of 3D renders represent oral region. Abbreviations: S, sinistral (left), D, dextral (right). Scale bars represent 5 cm in panels A,B, 1 cm in panels D-H.

### (c) Computed tomography

*Rhinopteraspis dunensis* NHMUK PV P 73217 (Figs 1,S1) was scanned using a Nikon Metrology XTH 225ST X-ray tomography instrument based in Bristol Palaeobiology, University of Bristol. Two scans were undertaken, each composed of two stacked scans. The first scan included the whole headshield at a voltage of 223 kV and a current of 193 μA, with 3141 projections, and with a 1 mm Sn filter, obtaining a dataset with 63.41 μm voxel resolution (Fig. 1A,B). The second scan targeted the oral region specifically, including the posterior third of the rostrum, the orbital, oral, and pineal parts of the headshield, and the anterior quarter of the dorsal and ventral discs, at a voltage of 180 kV and a current of 178 μA, with 3141 projections of 8 frames and 708 ms each, and with a 0.5 mm Cu filter, achieving a voxel resolution of 22.91 μm (Figs 1D-H,2).The resulting tomographic datasets were segmented in Mimics v.25 (materialise) to create 3D models. All 3D models were visualised in Blender 3.5 (blender.org).

### (d) Reconstruction and animation

3D models of the higher resolution scan set of *Rhinopteraspis* were imported into Blender for retrodeformation (Fig. 3). Use of Blender tools for retrodeformation followed the recommendations and techniques set out by Herbst et al. (52). Initial retrodeformation focused on the best preserved plates and those with clearly delineated articulations (53), including the rostral, orbital, supraoral and paraoral plates. After rearticulating and repairing deformation (see Supplementary Information), these elements provided a framework to delimit the dorsal and lateral extent of the oral plate array. The oral plates were aligned, maintaining their preserved order, within this delimited area by placing their dorsal tips close to the margin of the mouth and rotating the plates caudoventrally to match the angle of the surrounding plates. The postoral plates were then fitted to the proximal ends of the oral plates. The proximal ends of the oral plates were then readjusted into articulation with the sulcus preserved in the postoral plates (Fig. 2C), while maintaining alignment with the closely articulated lateral oral- and paraoral plates and each other. Finally the ventral disc was retrodeformed and articulated with the postoral plate. The reconstructed specimen was animated in Blender to simulate the movement of the oral plates Fig. S3). Empties (single geometry-less points that act as handles for object transformation without interfering in the render process) were placed between each oral plate and the postoral plate below, so that the local x-axis of each was aligned with the approximate outer boundary of the postoral plate. Each oral plate was then parented (=linked) to the empty below it. These empties were then animated to rotate around their local x-axes, causing the parented oral plate to also rotate around that axis.

**Figure 2.**
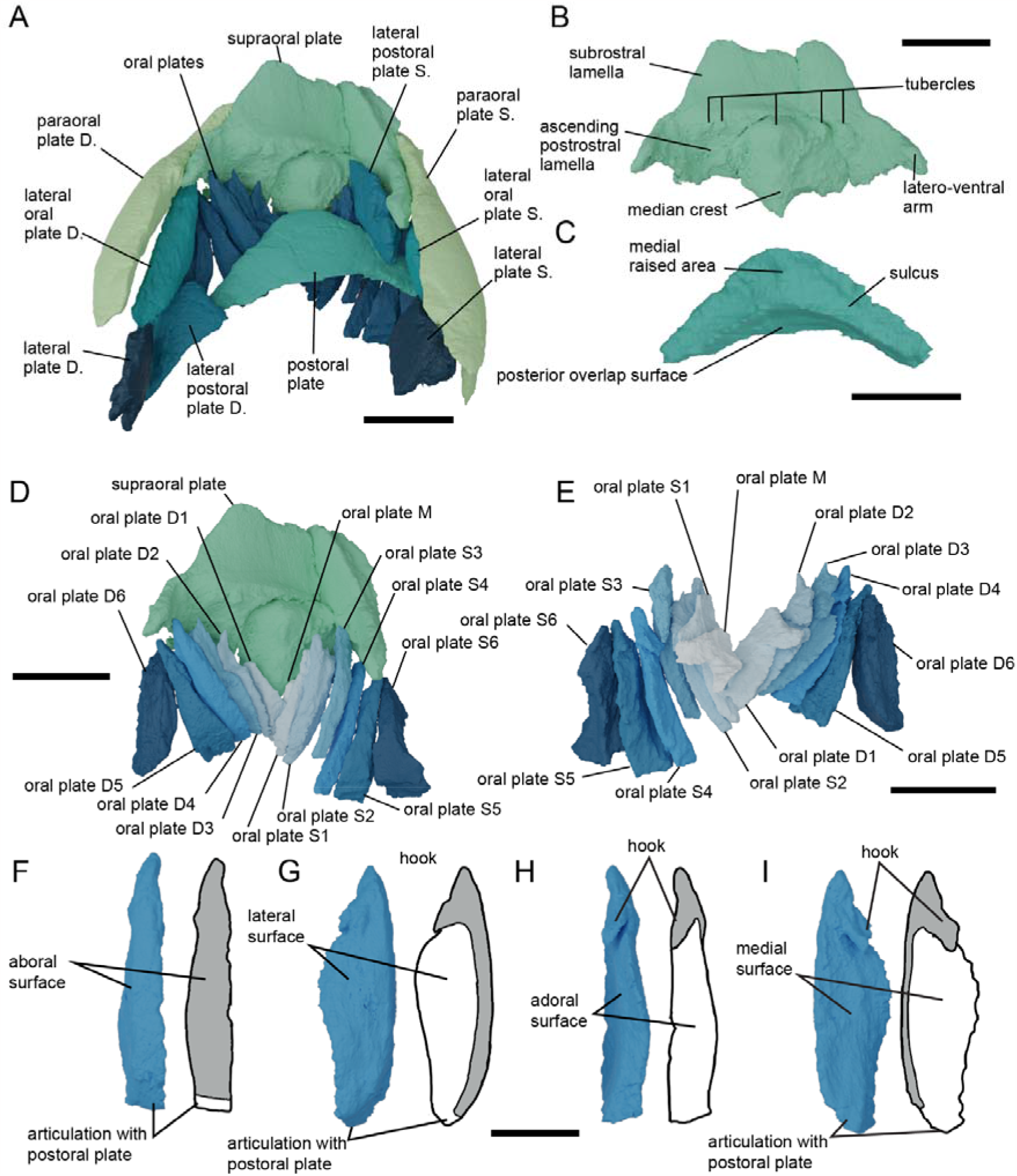
*Rhinopteraspis dunensis* NHMUK PV P 73217 oral region. A, oral apparatus and surrounding plates as preserved, in ventral view; B, ventral view of supraoral plate with three fragments rearticulated; C, postoral plate in dorsal view; D, oral apparatus as preserved with ventral plates removed; E, dorsal view of oral plates as preserved; F-I, oral plate R4 in aboral (F), lateral (G), adoral (H), and medial (I) views, alongside drawings depicting the inferred extent of dermal ornament in grey, based on comparison with isolated plates of Loricopteraspis dairydinglensis(20,40,58). Abbreviations: S, sinistral (left), D, dextral (right). Scale bars represent 1 cm in panels A-E, 0.5 cm in panels F-I.

**Figure 3.**
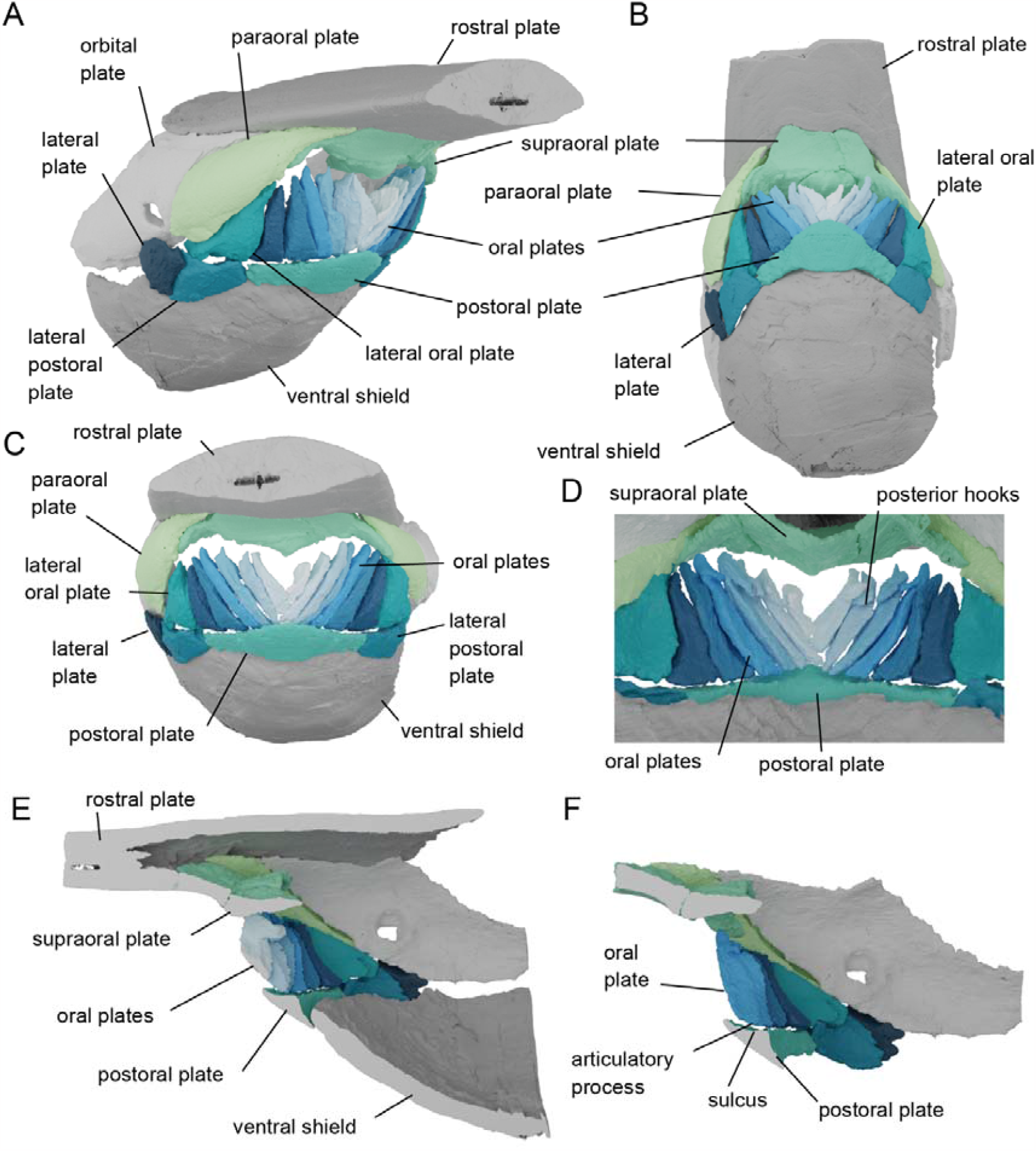
Reconstruction of *Rhinopteraspis dunensis* based on NHMUK PV P 73217, anterior section of rostrum not shown. A, rostrolateral view; B, ventral view; C, rostral view; D, caudal view of articulated oral plates; E, sagittal cross-section through centre of the head; F, close up cross-section lateral to E, showing junction between base of oral plate and the postoral plate.

## 3. Description

NHMUK PV P 73217 comprises an almost-complete three-dimensional specimen of *Rhinopteraspis *dunensis**, preserving the entire headshield and articulated body scales (Fig. 1A,B,S1). The specimen has been crushed laterally, with the oral region and ventral shield displaced rostro-dorsally (Fig. 1). Otherwise, the specimen is complete and individual oral elements appear to have maintained their original shape and relative location, as evidenced by their approximately symmetrical arrangement. The dermal skeletal anatomy of pteraspid heterostracans is well-characterised in numerous taxa (18,54) to which that of *Rhinopteraspis *dunensis** NHMUK PV P 73217 conforms (32,55,56). The headshield is composed of large dorsal and ventral shields separated by paired cornual and branchial plates, with a dorsal spine set into the posterior margin of the dorsal shield (Fig. 1A,B,S1). Anterior to the dorsal shield is an elongate rostrum that is separated from the dorsal shield by paired orbital plates (Figs 1A,B,S1). The anterior length of the rostrum is broken off from the rest of the specimen (Figs 1A,B,S1). The unpaired pineal plate is indistinguishable from the top of the orbital plates in the scan data.

The oral region is bordered dorsally by the supraoral plate, laterally by paired paraoral plates, and caudally by the postoral plate (Fig. 1). In previous descriptions of pteraspids, including *Rhinopteraspis*, the subrostral lamella and ascending postrostral lamella have been characterised as part of the rostral plate (51,57), although Friman & Bardenheuer described paired plates in this position they termed the ‘subrostral plates’ (32). In NHMUK PV P 73217 they comprise a separate structure, broken into three parts (postmortem), which we term the supraoral plate (Figs 1D,G,H, 2A,B). The supraoral plate is trapezoidal in shape, narrowing rostrally. A pronounced furrow runs around its lateral and rostral margins, which is overlapped by the paraoral,orbital, and rostral plates; the margins of the plate sweep ventro-caudally to form two latero-ventral arms (Fig. 2B). The rostral half of the ventral surface of the supraoral plate (the subrostral lamella) is surmounted by a superficial layer of ornament and made convex by a prominent median crest, aligned rostro-caudally (Fig. 2B). The caudalmost half (the ascending postrostral lamella) curves upwards into the mouth and lacks dentine ornament (as in *Rhinopteraspis cornubica* Tarlo, 1961, p. 373), the rostral border of this area presumably represents the position of the oral opening. This border is marked by a row of tubercles, comprising one large medial tubercle and two pairs of smaller tubercles on either side (Fig. 2B). The paired paraoral plates are elongate, taper rostrally, and overlie large rostro-lateral overlap surfaces on the orbital plates, rostral to the orbits (Figs 1D-F, 2A). We interpret the ‘olfactory grooves’ identified in *Rhinopteraspis cornubica* by Tarlo (1961, fig. 1) as the overlap surfaces between the paraoral plates and supraoral plate (Fig. 2A) (32).

The postoral plate is bow-shaped and originally symmetrical, although the right process is damaged (Fig. 2A,C). The ventral surface is smoothly convex. The inner surface has a caudal overlap surface that is concave to curve around the anterior rim of the ventral shield, and a dorsal surface that forms the ventral margin of the oral opening, bearing paired sulci for the oral plates that are interrupted medially by a raised area (Fig. 2C). Paired lateral plates and lateral postoral plates lie between the postoral plate and the orbital plates (Figs 1,2). The extensive overlap between the plates surrounding the mouth and the larger head shield plates strongly suggests that they comprised an integrated structural unit with little or no movement relative to each other.

The oral apparatus itself is composed of thirteen imbricated oral plates and one pair of lateral oral plates (Figs 1,2,S2). The aboral, mediolateral surfaces of each oral plate, as well at the lateral surfaces of the hook, are all faced with tuberculated ornament (Fig. 2F-I). However, the lateral and adoral surfaces of the oral plates, as well as the ventral side of the hook and the proximal end of the plates, are all unornamented, instead exhibiting a porous surface texture reflecting open vascular canals. The plates are arranged bilaterally about the midline into sinistral and dextral series. Although plate pairs that occur in equivalent positions on either side of the midline are similar, they do not exhibit mirror-image symmetry. In particular, the unpaired medial plate is not symmetrical but, rather, is continuous with the sinistral series (Fig. 2E,S2). Each oral plate has the same general morphology of a main limb with a rhomboidal cross-section, a distal hook (except for the most laterally placed plates), and a proximal articulation surface for the postoral plate (Figs 2,S2). There are six oral plates in the sinistral series (S1-6) and six in the dextral series (D1-6), each preserved inclined at varying degrees (maximum about 45 deg) to each other along their coronal axis. The more medially placed the plate, the more inclined it is along its long axis to provide a fit with the adjacent oral plate; the lateral and medial faces of the plates overlap and imbricate, inclined at increasing angles relative to the sagittal plane, from medial to lateral. The distal hooks of the oral plates curve adorally, while the proximal ends of the plates are notched, reflecting the ventral limit of the external dermal ornament, serving as articulation with the postoral plate (except for the medial plate M).

Although the oral plates have similar morphologies, individually they vary in relative proportion depending on their position within the apparatus (Figs 2,S2). The hook of the unpaired medial oral plate M is as long as the main limb. This plate is preserved overlying the lateral two oral plates D1 and S1. The lateral side of its proximal end is notched to fit with the left medial face of D1. Plate M curves dextrally such that it fits the curvature of the dextral face of the adjacent oral plate S1. The main limb of oral plates D1 and S1 are twice as long that of M, with a narrow base; similarly, it may not have articulated with the postoral plate, but the hooks in D1 and S1 are as large and very similar in shape. These plates fit around oral plates S2 and D2, which fit around S3 and D3, etc.; these pairs have narrow bases and slightly smaller hooks than S1 and D1. Plates S4 and D4 have a slightly broader base and proportionally shorter hook. This trend continues laterally, with increasingly shorter hooks and wider bases in S5 and D5. Finally S6 and D6 have no perceptible hook and a very broad base. The lateral edges of plates S6 and D6 is concave, fitting the medial edge of the lateral oral plates.

The lateral oral plates are morphologically distinct from the oral plates (Fig. 2,S2), approximately triangular, tapering rostrally with a curved medial margin that matches the lateral profile of the lateral-most oral plates (Figs 2A,S2). The better preserved sinistral lateral oral plate appears to have a distinct notch in its posterior side (Fig. S2A,C), although this is difficult to corroborate from the dextral lateral oral plate.

## 4. Reconstruction

The combined width of the bases of the oral plates, when aligned perpendicular to a sagittal plane, matches the length of the sulcus on the postoral plate (Fig. 3) and the complementary symmetry exhibited by adjacent plates indicates that they were capable of almost completely filling the width of the oral opening. In this position, the lateral-facing surface of each oral plate overlaps the adoral surface of its outer neighbour (Fig. 3). The lateral surfaces of the outermost oral plates fit closely with the lateral oral plates (Figs. 3C,D). M, S1 and D1 appear to lie on top of the neighbouring oral plates rather than contacting the postoral plate. When viewed caudally, this brings the hooks of the plates into alignment, forming a plane above the posterior unornamented surface of the plates (Fig. 3d). This reconstruction also suggests that the oral apparatus fits together such that the tops of the oral plates extend to the top of the mouth to almost meet the supraoral plate. The medial crest of the supraoral plate creates a convex dorsal margin of the mouth, and the increasing length of the oral plates in lateral positions means that their dorsal tips also follow this convex line. Thus, when fully closed, there would have remained a narrow “letterbox-shaped” opening (cf. (59)) into which projected the dorsal tips of the oral plates and the medial crest and associated tubercles of the dorsal oral plate. Indeed, the medial crest of the supraoral plate would have effectively divided the residual opening into two. There is no evidence that the plates intercalated with the tubercles demarcating the dorsal margin of the mouth at the rostral margin of the ascending postrostral lamella, Fig. 2B). When modelled to open around the axis of the sulcus on the postoral plate synchronously, the oral plates do not overlap as they rotate (Fig. S3). Instead, their placement along the curved axis of the postoral plate sulcus causes them to splay outwards (Fig. S3).

## 5. Discussion

Based upon our 3D reconstruction of the oral region in *Rhinopteraspis*, we are able to consider the oral plates as an integrated apparatus and test hypotheses of its form and function. The endoskeleton of heterostracans is completely unknown beyond what can be inferred from the dermal skeleton (60). Hence, we attempt to consider the constraints imposed by the dermal skeleton, without speculating as to endoskeletal structure.

### (i) Interpretation of the oral apparatus of *Rhinopteraspis*

Most previous hypotheses of function in heterostracan feeding assume significant movement and a degree of rotation of the oral apparatus, often following a cyclostome-like mode (19,22,24). In many of these the postoral plate(s) is assumed to move significantly, either inwards (24) or outwards (22,59). The large, curved overlap surface at a 45° angle between the postoral plate and the ventral plate in *Rhinopteraspis* (Fig. 2C) suggests that this movement would not have been possible and that the postoral plate was static during feeding. In contrast, the imbricated nature of the oral plates strongly suggests that they were mobile relative to the postoral plate. Any movement requires rotation around the point of their attachment to the postoral plate. The rhomboid cross-section and the complementary symmetry of the oral plates would have prevented them either moving independently, or moving in an entirely sagittal plane (Fig. S3). Rather, they must have moved as an integrated unit, splaying ventro-laterally as the plates rotated aborally on the sulcus of the postoral plate. The medial plates (M, S1, D1) that do not articulate with the postoral plate are the exceptions: their cross-sectional shape and ab/adoral overlap would have precluded their movement relative to the other plates.

The morphologies of individual oral plates are closely comparable to those observed in acid-prepared isolated oral plates of *Loricopteraspis dairydinglensis* recovered through acid digestion of limestone (20,40,58). The open vasculature on the lateral surfaces and adoral surfaces, as well as the ventral side of the hook and the proximal end of each plate, is significant. This indicates that these parts of the plates were embedded in soft tissue, the upper limit of which would have been the proximal surface of the large projecting hooks on the adoral side of the oral plates. The medial plates would have been supported entirely by this soft tissue which must have provided the basis of any movement of the plates and so may have included unmineralized cartilage, muscles or tendons.

Our reconstruction indicates that, even when the mouth was fully closed by occlusion of the oral plates, there would have remained a short but wide gape that was effectively divided in two by the median crest of the supraoral plate. The associated tubercles of the supraoral plate and tips of the oral plate would have projected into this space, further occluding it. This oral morphology precludes interpretation as a hagfish-like prenasal sinus (26,27) because the opening is too small. There is also no evidence for separate upper oral plates as inferred by Stensio (26,27). Assuming the presence of forward pointing denticles on the lateral surfaces of the oral plate hooks, as in *Loricopteraspis* and *Pteraspis* (20,58), these structures would not have bordered the oral opening but, rather, the junction between each plate and its neighbour, linked by soft tissue. Rather than being involved in food capture or processing, this ornamentation may have helped to prevent particles from becoming lodged in the spaces between the plates (58) and their associated soft tissue, preventing fouling of the oral apparatus (cf. Hamann and Blanke (1)). Any movement seems unlikely to have been great in magnitude due to the weak joint between the oral plates and the postoral plate, as well as the suspension of the small median oral plates in soft tissue. A structural analogue in a living vertebrate for heterostracan oral plates might be the branchiostegal plates in osteichthyans, which support the branchiostegal membrane, and make limited, coordinated movements to aid the suction pump (61).

The oral apparatus of *Rhinopteraspis* could only have been moved from the adoral side with rostro-caudal movements, as the aboral surfaces of the oral plates lack any kind of attachment surfaces. This is inconsistent with a gnathostome-like organisation of paired mandibular adductor muscles. However, living cyclostomes both operate the oral apparatus by moving cartilages rostrally and caudally along the floor of the pharynx. In hagfishes, keratinous toothlets are mounted on a cartilaginous dental plate that is pulled anteriorly along a basal plate to evert the lingual apparatus, and posteriorly to return it to resting position (5,6,8). In lampreys, a medial piston cartilage is protracted rostrally, moving a medial apical cartilage, the action of which brings keratinous toothlets in front of the apical cartilage into contact with other toothlets in a rasping action (7,62,63). A medial groove along the visceral surface of the ventral shield in some pteraspids has been cited as evidence for a cyclostome-like medial structure of the oral mechanism (60).

### (ii) Comparison of oral anatomy with other heterostracans

The anatomy of the oral apparatus of *Rhinopteraspis* is comparable to that of other pteraspids. Individually, the shape of the oral plates conforms closely to the morphologies of isolated plates acid prepared from an articulated specimen of *Loricopteraspis dairydinglensis* ((40), pl. 37; (58)). The arrangement of these plates is similar to articulated specimens of *Protopteraspis* (19) and *Errivaspis* (22). Importantly, the posterior alignment of the hooks in *Rhinopteraspis* (Fig. 3) can also be seen in other pteraspids where the adoral side of an articulated oral apparatus is visible: Mylopteraspidella (Stensiö, (27), Fig. 44, p. 197) and *Protopteraspis* (19), as well as in *Errivaspis* (White, (22)figs 41-44). Together, these indicate that our interpretation of the feeding apparatus of *Rhinopteraspis* is more broadly applicable within pteraspids. However, varied body shapes and positions of the oral opening indicate some diversity in feeding ecology (e.g. *Doryaspis* (64), *Drepanaspis* (65)).

*Athenaegis*, the oldest articulated heterostracan (66), is often assumed to be an outgroup to all other macromeric heterostracans (21,50,67,68), and its oral apparatus appears broadly anatomically consistent with *Rhinopteraspis*. By contrast, cyathaspids, a likely paraphyletic grade of heterostracans (21), have arrangements of plates that are difficult to rationalise with the oral apparatuses of *Athenaegis* and pteraspids (e.g. *Anglaspis* (37), *Poraspis* (69), *Capitaspis* (70), and Allocryptaspis (29)). In amphiaspids, the oral aperture occurs at the end of a tube in a single, fused headshield (71). Meanwhile the oral apparatuses of the “tessellate heterostracan” taxa are completely unknown (34) and what is known of their anatomy is difficult to reconcile with the pteraspids, including *Rhinopteraspis*. Thus, while the oral apparatus of *Rhinopteraspis* may be representative of pteraspids, *Athenaegis* and, by implication, macromeric heterostracans primitively, it may not be representative of heterostracans more generally, which may have exhibited greater diversity in terms of feeding ecology. Further investigation is required to assess the ubiquity of suspension-feeding within the group, and the position in the water column at which they fed.

### (iii) Implications for feeding in pteraspids and ancestral heterostracans

Our articulated oral apparatus model can be used to consider the various feeding strategies that have been proposed for pteraspid heterostracans. These strategies can be roughly divided into macrophagy, predation, deposit/detritus feeding, and microphagy or suspension feeding (72).

Hypotheses of macrophagous and predatory heterostracan feeding are based on analogy between heterostracan oral plates and the gnathostome mandible (e.g. (19)), or more rarely the hagfish oral apparatus (e.g. (24)). However, the oral apparatus in *Rhinopteraspis* is poorly constructed for biting or grasping. Although the ascending lamella bears tubercles that have been interpreted as the upper ‘jaw’ analogue in such a scenario (57), they do not occlude with the tips of the oral plates. Moreover, the angle of approach of the plates to the preoral plate during closing is oblique, with a low mechanical advantage, and would have been poorly suited to generating force. Finally, the oral plates themselves would not have formed a firm biting surface, with a poorly developed joint with the postoral plate even in the broad based lateral oral plates (e.g. plates S/D4-6), and the medial oral plates (e.g. plates M, S1, D1) suspended in soft tissue or with very narrow-based articulations with the postoral plate. Scenarios that envisage see the oral plates as elements analogous to hagfish-like toothlets (24) can be similarly rejected because the junction between the proximal tips of the oral plates and the sulcus in the postoral plate, and the fixed position of the postoral plate, would all have acted to dramatically restrict the movement of the oral plates.

Deposit or detritus feeding interpretations typically envisages the oral apparatus as a ‘scoop’ that would have been used to acquire sediment from the bottom of the water body (22). This requires the plates to evert substantially from their resting position, with some reconstructions also require movement of the postoral plate (22 figs 49,50). We can reject this based on our reconstruction of *Rhinopteraspis*, where significant movement of the oral plates is limited by the joint between the postoral and ventral plates. However, we cannot rule out the burial of the snout/mouth in the substrate as a means of deposit feeding. The presence or absence of wear, which is often used as a line of evidence in discussion of deposit feeding (20,73), are not visible in *Rhinopteraspis* at the resolution of our scan data. We note though that the long-snout in pteraspids such as *Rhinopteraspis*, along with the strongly convex ventral abdomen, would have restricted the ability of the animal to place its mouth in contact with the substrates.

In either suspension feeding or deposit feeding modes, rostral rotation of the oral plates would have resulted in a greater area of gape, increasing intake. The occluded oral apparatus leaves a restricted opening with digitate margins defined by the tips of the oral plates and the tubercles of the dorsal oral plate that would have served to prevent fouling (cf. (1,58)) while facilitating inflow in a manner analogous to straining functions seen in animals as diverse as flamingos, brachiopods, oysters and gastropods (74). It is possible that the exposed tips of the oral plates in *Rhinopteraspis* acted as an analogous structure along with the tubercles on the supraoral plate. The limited movement acted to control the entry of larger particles into the mouth and may also have served to provide a means to expel larger particles trapped between the plates (1).

### (iv) Implications for early vertebrate evolution

The New Head and New Mouth hypotheses (9,12) have argued for a long term shift towards increasingly active food acquisition, from the filter-feeding of invertebrate chordates to macrophagous predation in living jawed vertebrates. The feeding ecology of the extinct heterostracans is key since this group represents one of the earliest diverging members of the gnathostome stem-lineage. A diversity of feeding ecologies have been attributed to heterostracans, from macrophagous predation, through scavenging and herbivory, to deposit and filter feeding. Our three-dimensional reconstruction of the feeding apparatus of *Rhinopteraspis dunensis* suggests its finger-like oral plates were limited to only a small degree of concerted rotation, precluding all proposed feeding strategies bar suspension feeding or deposit feeding by burial of the mouth in the substrate. Comparison to other heterostracans suggests that while this condition may be unrepresentative of the clade generally, it is representative of pteraspids and the oldest and earliest well-known heterostracan; as such it may be plesiomorphic for the clade. Given macrophagy in earlier diverging lampreys, hagfish and conodonts (72), it is clear that early vertebrates and stem-gnathostomes had already established a diversity of feeding ecologies, long before the origin of jaws. This finding is consistent with recent challenges to the predictions of New Head Hypothesis, demonstrating that vertebrate innovations and elaborations cannot be characterised by a directional trend towards increasingly active food acquisition (75–77), but, rather, increasing ecological diversity.

## 6. Conclusion

A lack of knowledge of the 3D anatomy of early vertebrate feeding apparatuses has obscured their feeding ecology, and in turn hindered testing macroecological scenarios seeking to explain early vertebrate evolution. Using X-Ray microtomography and computed tomography we have reconstructed the oral apparatus of the pteraspid heterostracan *Rhinopteraspis dunensis*. The oral apparatus in *Rhinopteraspis* is composed of one medial and six pairs of bilaterally arranged oral plates, plus a pair of lateral oral plates. Inferred articulation of these plates indicates that range of motion was limited; the oral plates could only move in concert and could not rotate far. When occluded, the oral plates left a short wide gape to the mouth, closed partially by a convex crest extending from the supraoral plate and the distal tips of the oral plates. The reconstructed anatomy precludes all proposed feeding modes bar suspension or deposit feeding in *Rhinopteraspis*. Heterostracans more generally show a far wider range of oral anatomies and body shapes than in pteraspids (29), and a diversity of feeding ecologies was established early in vertebrate evolution, long before the origin of jaws. Given the existence of contemporary macrophagous vertebrate predators and scavengers, the presence of suspension feeding heterostracans is incompatible with the directional trend towards increasingly active food acquisition envisaged by the New Head hypotheses.

## Acknowledgements

Many thanks to Liz Martin-Silverstone (University of Bristol) for help with CT scanning. Thanks also to Emma Bernard (NHMUK) for specimen access.

## Data Availability

On publication all CT data and 3D models upon which this study is based will be made publicly and freely available.

## Supplementary Material Captions

**Supplementary Figure 1.**
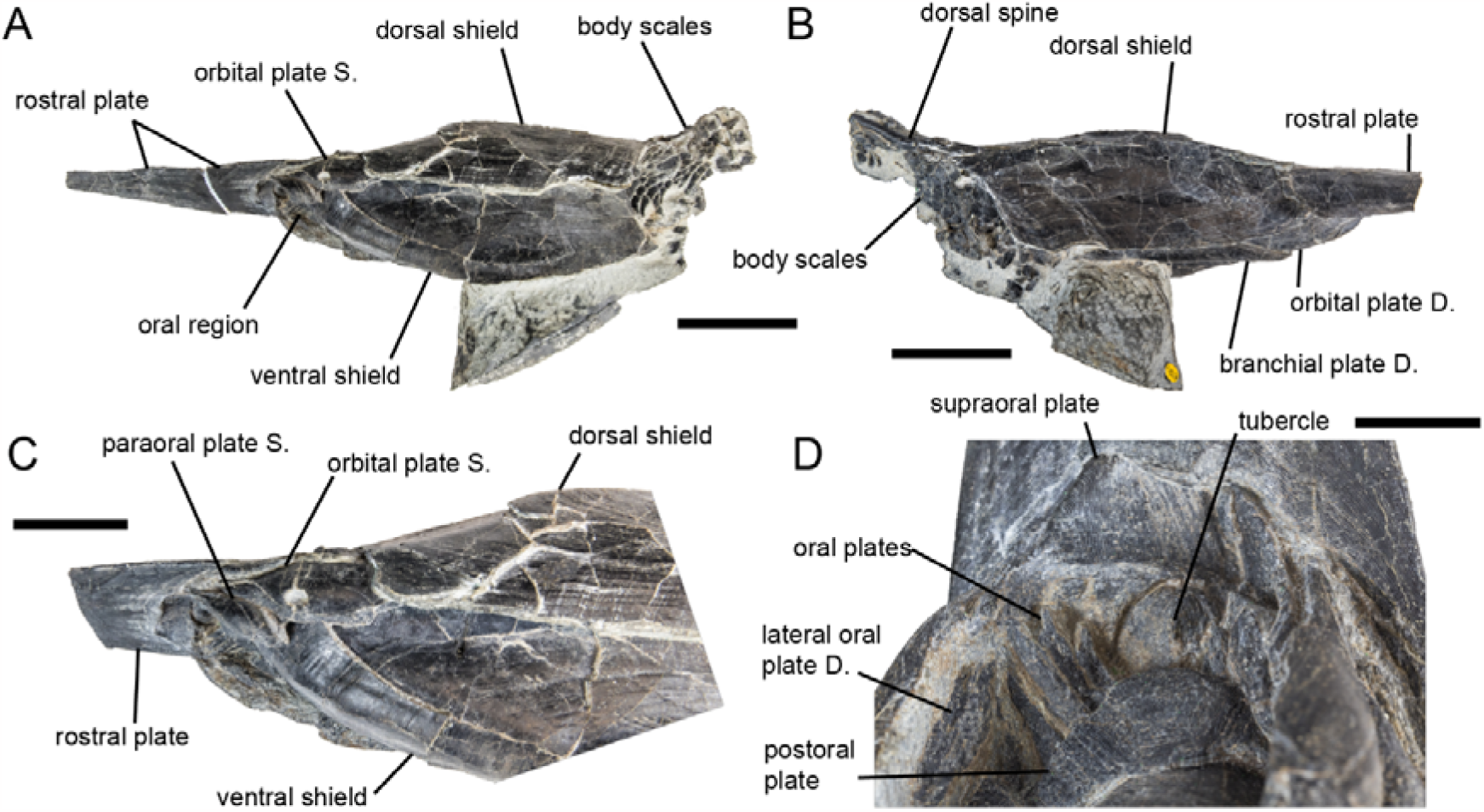
*Rhinopteraspis dunensis* NHMUK PV P 73217. A-D, photographs of complete specimen in (A) left lateral view with full length of preserved rostrum, (B) right lateral view, (C) close-up of front of head in left lateral view, and (D) close-up of oral region in ventral view; E-G, Abbreviations: S, sinistral (left), D, dextral (right). Scale bars represent 5 cm in panels A,B, 2.5 cm in panel C, 1 cm in panel D.

**Supplementary Figure 2.**
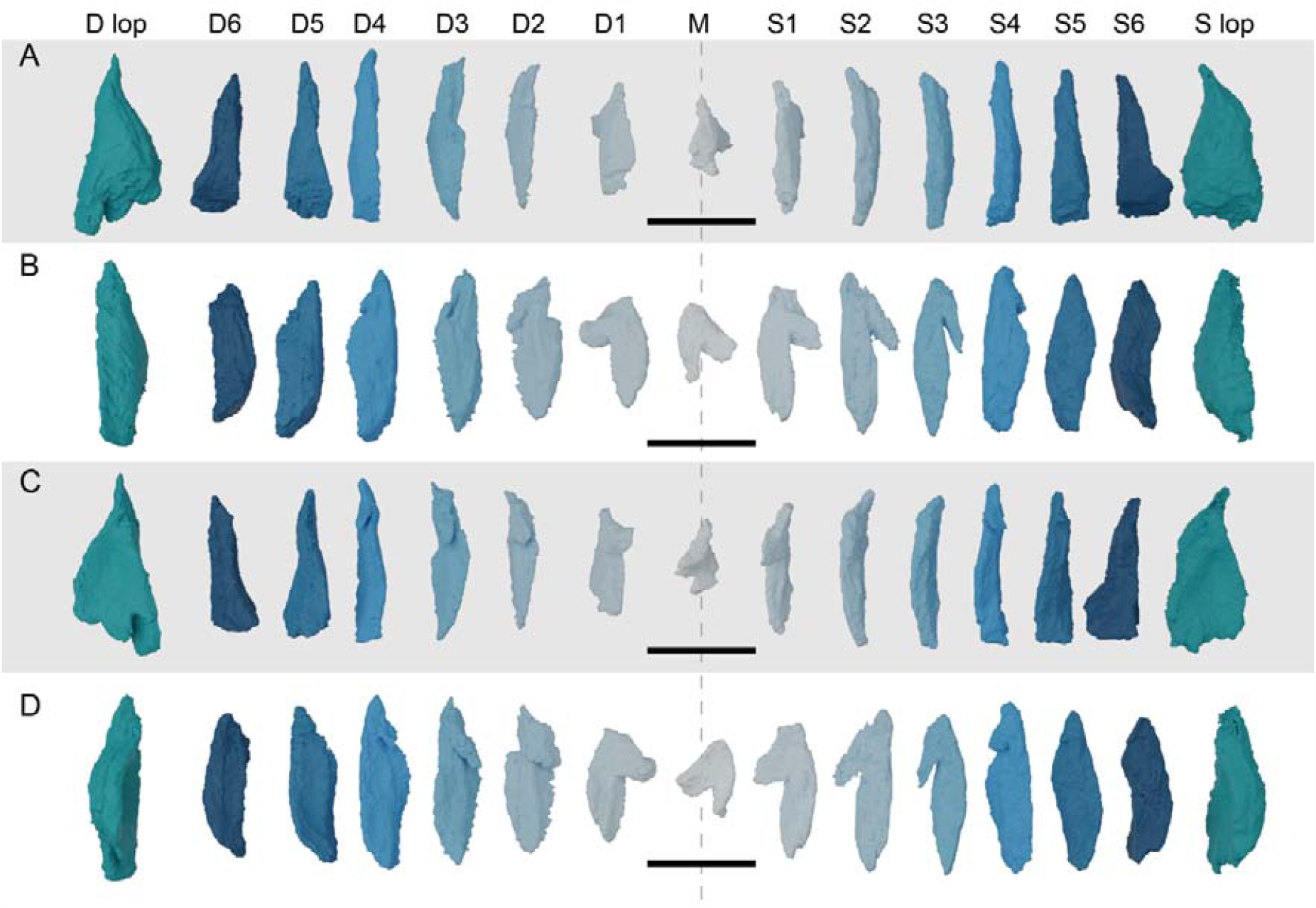
*Rhinopteraspis dunensis* NHMUK PV P 73217 oral and lateral oral plates in aboral (a), lateral (b), adoral (c), and medial (d) views. Dotted line intersects unpaired oral plate M. Abbreviations: lop, lateral oral plate, S, sinistral (left), D, dextral (right). Scale bar represents 1 cm.

**Supplementary Figure 3.**
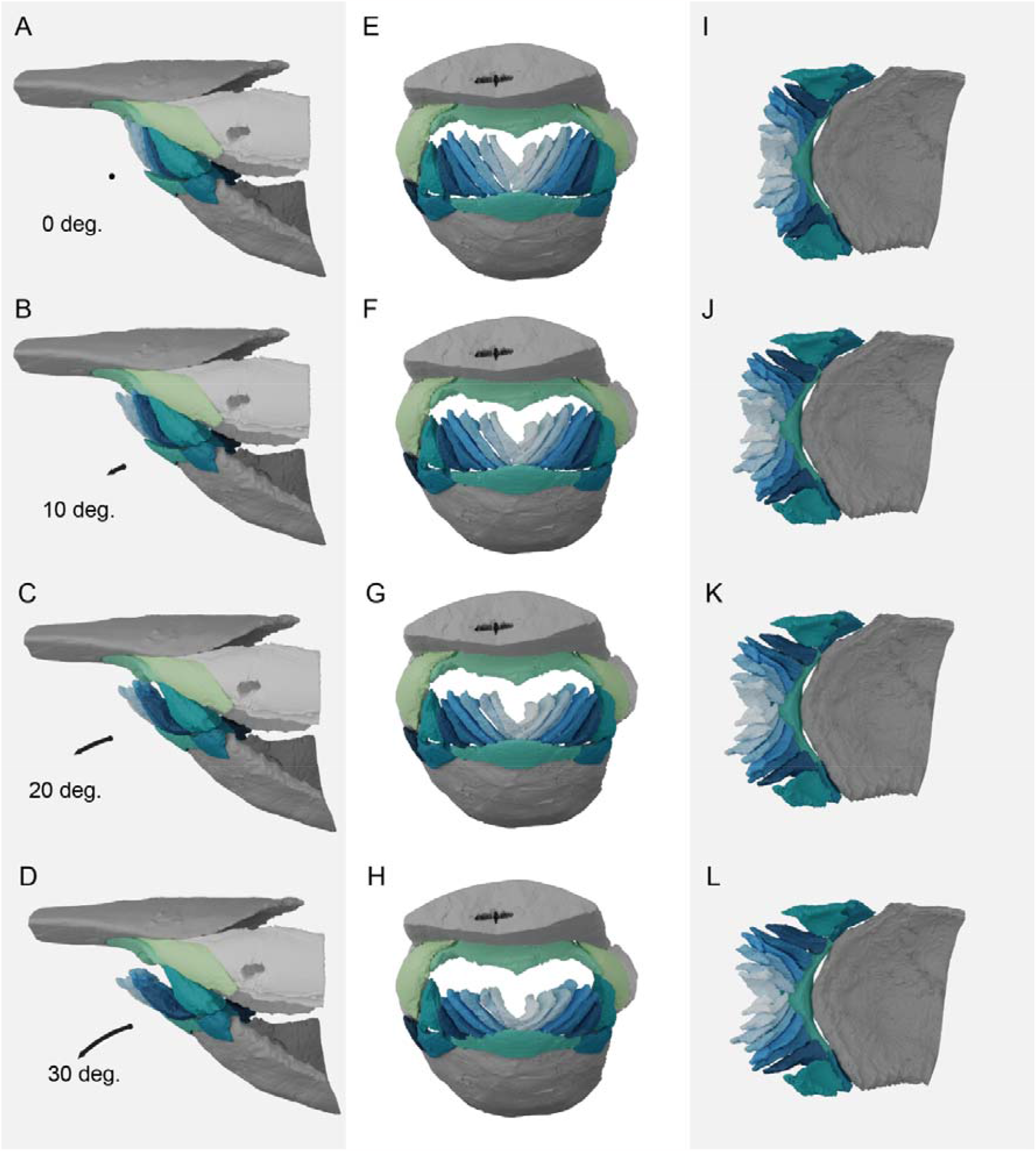
Reconstruction of *Rhinopteraspis dunensis* based on NHMUK PV P 73217 with oral plates animated to open to 30 degrees. A-D, lateral view; E-H, anterior view; I-L, dorsal view (with surrounding plates removed). A,E,I, resting position; B,F,J, open to 10 degrees; C,G,K, open to 20 degrees; D,H,L, open to 30 degrees.

